# The Toll pathway mediates *Drosophila* resilience to *Aspergillus* mycotoxins through specific Bomanins

**DOI:** 10.1101/2022.08.18.504437

**Authors:** Rui Xu, Yanyan Lou, Antonin Tidu, Philippe Bulet, Thorsten Heinekamp, Franck Martin, Axel Brakhage, Zi Li, Samuel Liégeois, Dominique Ferrandon

## Abstract

Host defense against infections encompasses resistance, which targets microorganisms for neutralization or elimination, and resilience/disease tolerance, which allows the host to withstand/tolerate pathogens and repair damages. In *Drosophila*, the Toll signaling pathway is thought to mediate resistance against fungal infections by regulating the secretion of antimicrobial peptides, potentially including Bomanins. We found that *Aspergillus fumigatus* kills *Drosophila* Toll pathway mutants without invasion because its dissemination is blocked by melanization, suggesting a role for Toll in host defense distinct from resistance. We report that mutants affecting the Toll pathway or the 55C *Bomanin* locus were susceptible to the injection of two *Aspergillus* mycotoxins, restrictocin or verruculogen. The vulnerability of 55C deletion mutants to these mycotoxins was rescued by the overexpression of Bomanins specific to each challenge. Mechanistically, flies in which *BomS6* was expressed in the nervous system exhibited an enhanced recovery from the tremors induced by injected verruculogen and displayed improved survival. Thus, innate immunity also protects the host against the action of microbial toxins through secreted peptides and thereby increase its resilience to infection.

## Introduction

The outcome of an infection depends on the interactions between a host and a pathogen with its armory of multifarious virulence factors. In the case of fungal pathogens, several hundred potential virulence factors are known to be secreted (Gao et al., 2011; Lebrigand et al., 2016).The host confronts the invading microorganism through the multiple arms of its immune system as well as varied strategies that counteract the effect of toxins and more generally withstand and repair damages inflicted directly by the pathogen or indirectly by the host through its own immune response (Ferrandon, 2013; Medzhitov et al., 2012; Soares et al., 2017). Fungal infections represent a widespread major health threat worldwide affecting more than 150 million patients and cause directly or indirectly at least one and a half million of deaths each year (Bongomin et al., 2017; Rodrigues and Nosanchuk, 2020). Our current understanding of fungal infections relies on the study of the host’s innate and adaptive immune responses and in parallel on investigations of fungal virulence factors (Scharf et al., 2014; van de Veerdonk et al., 2017). *A. fumigatus* can synthesize and secrete a vast array of toxins and secondary metabolites, the *in vivo* functions of which are just starting to be deciphered (Frisvad et al., 2009; Macheleidt et al., 2016; Raffa and Keller, 2019). Whereas some fungal virulence factors allow *A. fumigatus* to elude detection by the immune system, a few mycotoxins such as gliotoxin or fumagillin are known to interfere with immune signaling and help neutralize immune cell functions (Cramer et al., 2006; Konig et al., 2019; Kupfahl et al., 2006). However, it is currently poorly known whether the innate immune system is able to detect and counteract the actions of mycotoxins through specific effectors.

*Drosophila melanogaster* represents a genetically amenable model system that is well-suited to study infections and innate immunity as there is no vertebrate-like adaptive immunity. Its innate immune system comprises several arms, including a cellular response mediated by plasmatocytes in the adult, melanization, which depends on the cleavage of pro-phenol oxidases by Hayan, and the humoral systemic immune response (Lemaitre and Hoffmann, 2007; Liegeois and Ferrandon, 2022). A landmark study published 25 years ago established the central role of the Toll pathway in mediating the humoral immune response against fungal infections, as represented by *A. fumigatus* (Lemaitre et al., 1996). This observation has since been extended to a variety of other filamentous fungi or pathogenic yeast infections and also to several Gram-positive bacterial infections. The current paradigm is that upon sensing infections, a MyD88 adaptor-dependent intracellular signaling pathway gets activated by the binding of the processed Spätzle cytokine to the Toll receptor and stimulates the transcription of effector genes that mediate its role in host defense, such as antimicrobial peptides (AMPs). Genes coding antifungal peptides such as Drosomycin, Metchnikowin, and Daisho are Toll-regulated AMPs active on filamentous fungi (Cohen et al., 2020; Fehlbaum et al., 1995; Levashina et al., 1995). However, in contrast to the other NF-κB signaling pathway that mediates host defense against Gram-negative bacteria, Immune deficiency (IMD), it is less clear whether Toll-dependent AMPs provide the bulk of the protection against Gram-positive or fungal infections. Indeed, the deletion of a locus encoding ten Toll-dependent secreted peptides at the 55C locus known as Bomanins largely phenocopies the *Toll* mutant phenotype (Clemmons et al., 2015; Uttenweiler-Joseph et al., 1998). This suggests that these peptides are somehow involved in mediating the defenses resulting from Toll pathway activation, which regulates the expression of more than 200 immune-responsive genes (De Gregorio et al., 2002). A majority of Bomanins at the 55C locus are short (BomS), the secreted form of which essentially contains a single domain characteristic of this family of peptides. Other members include a tail after the Bomanin domain (BomT) whereas bicipital members are characterized by the inclusion of two omains separated by a linker domain (BomBc). Although a recent study suggests that some BomS are likely active against *Candida glabrata* and can function somewhat redundantly (Lindsay et al., 2018), the exact function of most Bomanins in host defense remains uncertain.

How exactly *Drosophila* host defense confronts *A. fumigatus* and more generally fungal virulence factors remains unknown despite our knowledge of the generic role of the Toll pathway in antifungal defense. Here, we revisit *A. fumigatus* infections obtained by injecting a limited number of conidia into the thorax and find that the fungus is unable to invade flies, including Toll pathway *MyD88* mutants, due to melanization, a distinct host defense, which is mediated by the Hayan protease and the PPO2 phenol oxidase. Our data suggest that Toll pathway immuno-deficient flies succumb to *A. fumigatus* secreted toxins, some of which target the nervous system. We report here that Toll mediates resilience to particular mycotoxins through specific Bomanins that do not function as classical AMPs in this setting but neutralize the effects of these fungal virulence factors. Our data illustrate that evolution has selected a specialized defense partially mediated by secreted peptides that elude or counteract the attack by mycotoxins.

## Results

### Defense against *A. fumigatus* depends on the Toll pathway independently from its role in controlling AMP expression

Homozygous or hemizygous *MyD88* but not wild-type flies were highly sensitive to an *A. fumigatus* challenge with various strains and succumbed to as few as five injected conidia (Fig. 1*A*; Fig. EV1*A-C*). Mutations in the *Drosophila* Toll pathway genes *spätzle* (*spz*) and *Toll* (*Tl*) led to an *A. fumigatus* susceptibility phenotype similar to that of *MyD88* (Fig. EV1*D-E*).

**Figure 1.**
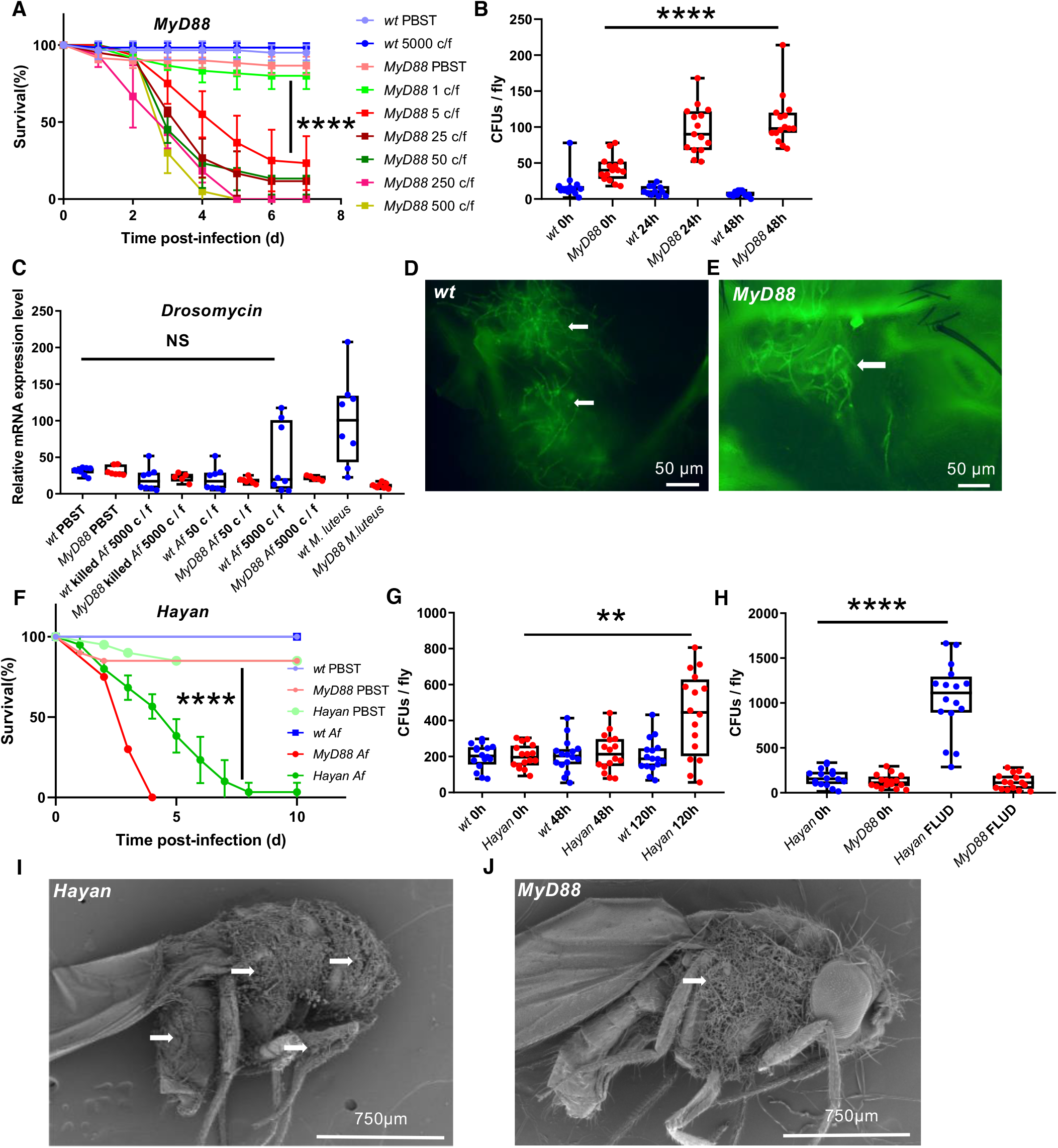
Toll pathway mutants succumb to *A. fumigatus* infection even though it is not required to limit the proliferation and dissemination of the pathogen, an immune function mediated by melanization. (*A*) survival curves of *MyD88* flies injected with different doses of *A. fumigatus* conidia (c/f: conidia injected per fly; mean and standard deviation of the survival of triplicates of 20 flies each are displayed). (*B*) fungal loads of single *MyD88* mutant and wild-type flies (50 conidia injected per fly). (*C*) expression level of *Drosomycin* induced by different doses of injected *A. fumigatus* conidia measured by RT-qPCR; *M. luteus* (OD=10) represents the positive control. (*D-E*) GFP-labeled *A. fumigatus* (50 conidia per fly) injected in wild-type (*D*) or *MyD88* mutant (*E*) flies form hyphae in the thorax of the flies (arrows). (*F-H*) *Hayan* flies are susceptible to *A. fumigatus*. Survival (*F*), time course of fungal loads of single *Hayan* mutant flies (500 conidia per fly) (*G*), and fungal load upon death (500 conidia per fly) (*H*) of *Hayan* mutant flies; *Hayan* 0h *vs*. 120h, p=0.004. (*I-J*) *A. fumigatus* hyphae extrusion (arrows) from *Hayan* (*I*) and *MyD88* (*J*) mutant cadavers; scanning electron micrographs. *B,C,G,H*: the middle bar of box plots represents the median and the upper and lower limits of boxes indicate respectively the first and third quartiles; the whiskers define the minima and maxima; data were analyzed using the Mann-Whitney statistical test. Survival curves were analyzed using the LogRank test. NS: not significant.

Unexpectedly and in contrast to other relevant microbial infections in Toll pathway mutants (Alarco et al., 2004, Wang, W et al., in preparation; Apidianakis et al., 2004; Duneau et al., 2017; Quintin et al., 2013) (Wang, W *et al*., in preparation), the fungal burden did not increase much in *MyD88* flies challenged with 50 conidia (Fig. 1*B*) or even upon the injection of 5,000 conidia (Fig. EV1*F*). The lack of growth of *A. fumigatus* in *MyD88* flies was confirmed by measuring the fungal load upon death (FLUD; Fig. EV1*G*). Monitoring a GFP-expressing *A. fumigatus* strain revealed the formation of mycelia only next to the injection site of 50 conidia in both wild-type and *MyD88* flies (Fig. 1*D-E*; Fig. EV1*H*). Puzzlingly, the injection of a higher number of conidia led to the formation of fewer hyphae (Fig. EV1*I*). To exclude the possibility that death might be caused by another microorganism, possibly deriving from the microbiota, we confirmed the sensitivity of *MyD88* mutant flies to *A. fumigatus* challenge on antibiotics-treated flies as well as on axenic flies (Fig. EV1*J-L*). We conclude that *MyD88* flies succumb to a low *A. fumigatus* burden (lower than 200 *A. fumigatus* colony-forming units [cfu] at death; Fig. EV1*G*).

A septic injury with the Gram-positive bacterium *Micrococcus luteus* induces the expression of *Drosomycin* and all 55C *Bomanin* genes, *BomS4*-excepted (Fig. EV2*A-B*). In contrast, the injection of even high doses of live or killed conidia did not induce the expression of *Drosomycin* steady-state transcripts measured by conventional RTqPCR as read-out (Fig. 1*C*). Only a mild induction of *Drosomycin* and the small secreted peptide-encoding gene *BomS1* were detected using digital PCR (RTdPCR) in wild-type flies challenged with 5,000 conidia (Fig. EV2*C-D*), which was confirmed by mass-spectrometry detection of the induction of some BomS peptides but not Drosomycin in collected hemolymph (Fig. EV2*E-F*). *A. fumigatus* infection thus induces weakly at the transcriptional level the expression of classical Toll pathway activation read-outs such as *BomS1* or *Drosomycin*. Surprisingly, only the short Bomanins and one Daisho peptide (DIM4) were reliably detected in the hemolymph. Their levels in the hemolymph was rather independent of the size of the inoculum (Fig. EV2*G*), in keeping with the relatively stable fungal load measured (Fig. EV1*F*). The expression of these peptides in the hemolymph tended to actually decrease upon injection of an inoculum> 1,000 conidia, possibly correlating with the decreased formation of hyphae and possibly higher levels of gliotoxin.

Thus, the SPZ/Toll/MyD88 cassette is required for host defense against *A. fumigatus* infections, even though this pathogen only mildly stimulates the Toll pathway and hardly its major read-out, *Drosomycin* steady-state transcript levels and its encoded peptide.

### Drosophila *melanization curbs* A. fumigatus *invasion*

As melanization is a host defense of insects effective against fungal infections, we tested *Hayan* mutant flies defective for this arm of innate immunity (Nam et al., 2012). These mutant flies were sensitive to *A. fumigatus* infection but less susceptible than *MyD88* mutant flies (Fig. 1*F*). In contrast to *MyD88*, the fungal burden was increased in these mutants (Fig. 1*G-H*). Further, the melanization response was dependent on Prophenoloxidase 2 (PPO2) but not PPO1 nor on the Sp7 protease (Fig. EV3*A-D*), in contrast to a previous study with *Enterococcus faecalis* (Dudzic et al., 2019). Interestingly, *A. fumigatus* disseminated throughout the body in *Hayan* mutants but was restricted to the thorax in *MyD88* flies (Fig. EV3*E*). In keeping with these results, the fungus erupted in cadavers from all three tagmata, including the legs, of *Hayan* mutants (Fig. 1*I*). In contrast, the fungus only broke through the cuticle in the thorax where it had been injected in *MyD88, Toll* or *spz* mutants (Fig. 1*J*, Fig. EV3*F*). Of note, the fungus did not erupt from infected wild-type flies killed mechanically. We conclude that melanization limits the proliferation and the dissemination of *A. fumigatus* injected into wild-type flies yet does not eradicate it at the injection site, where a melanization plug forms.

We also tested the contribution of the cellular immune response either by challenging *eater* mutant flies lacking a major phagocytosis receptor, pre-saturating the phagocytic apparatus by injection of latex beads, or by genetically ablating hemocytes. In each case, no enhanced sensitivity to *A. fumigatus* infection was observed (Fig. EV4).

### A. fumigatus *secondary metabolism is required for its virulence in* Drosophila

The finding that *A. fumigatus* killed *MyD88* immuno-deficient flies with a low fungal burden and limited dissemination in conjunction with the observation of the, at most, modest induction of the Toll pathway suggested that Toll pathway mutants could be sensitive to some of the many diffusible mycotoxins known to be secreted by this fungus (Frisvad et al., 2009).

We first tested this hypothesis using an *A. fumigatus* mutant strain lacking the phosphopantetheinyl transferase (*pptA*) gene required for the biosynthesis of all secondary metabolites, including most mycotoxins (Johns et al., 2017). This *A. fumigatus* mutant strain was not virulent when its conidia were injected into *MyD88* flies (Fig. 2*A*), even though they did manage to form a limited mycelium at the injection site (Fig. 2*C-D*); the fungal burden of the Δ*pptA* mutant was somewhat reduced after 48h (Fig. 2*B*). These findings indicated that one or several mycotoxins are responsible for the observed phenotypes. However, gliotoxin was not required to kill *MyD88* mutant flies as a gliotoxin deletion mutant, *Δglp*, was still as virulent as wild-type *A. fumigatus* (Fig. 2*E*). As expected from the analysis of the gliotoxin mutant strain, the injection of commercially-available gliotoxin killed wild-type and *MyD88* flies at a similar rate, but only when injected at sufficiently high concentrations. By contrast, fumagillin and helvolic acid did not kill wild-type or *MyD88* flies at the tested concentrations (Fig. 2*F-H*).

**Figure 2.**
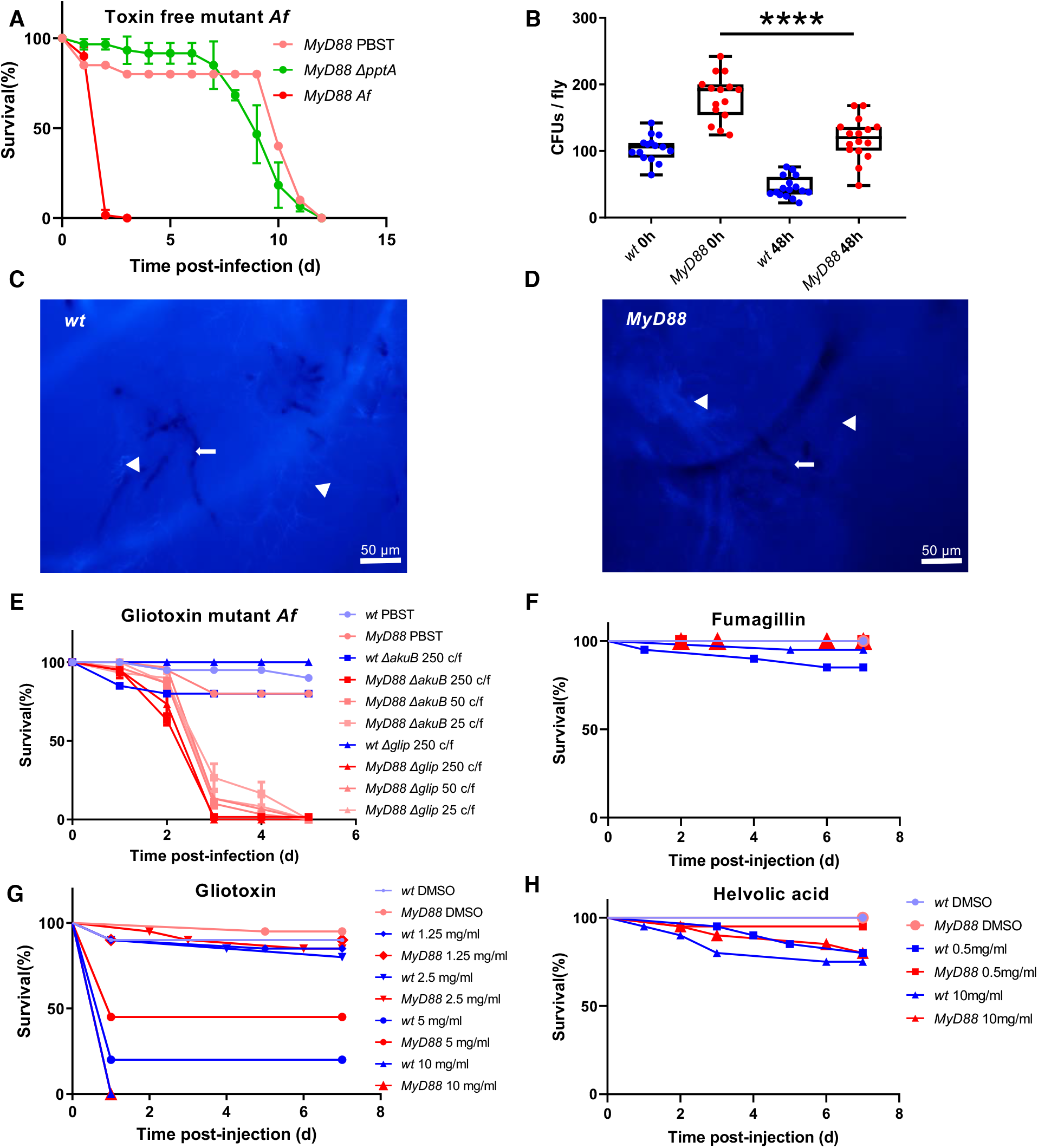
Secondary metabolism is critical for the virulence of *A. fumigatus* in *Drosophila MyD88* immunodeficient flies. (*A*) survival of *MyD88* flies injected with *Δ1pptA* and wild-type (*Af*) *A. fumigatus* conidia; Mean and standard deviation of the survival of triplicates of each 20 flies are displayed. (*B*) fungal loads of single flies after the injection of 500 *Δ1pptA* conidia. (*C-D*) hyphae of *Δ1pptA A. fumigatus* observed in thorax of wild-type and *MyD88* flies (arrow) after UVITEX negative staining, air sacs and tracheae are stained by UVITEX (arrowheads). (*E*) dose response of *MyD88* flies after *Δ1gliP* (gliotoxin) mutant or wild-type [*Δ1akuB*] *A. fumigatus* infection mean and standard deviation of the survival of triplicates of each 20 flies are displayed; wild-type flies are used as a control for the dose of 250 conidia. (*F-H*) dose response of *MyD88* and wild-type flies after gliotoxin (*F*), fumagillin (*G*), and helvolic acid (*H*) injection at the indicated concentrations (n=20 per condition). *B*: the middle bar of box plots represents the median and the upper and lower limits of boxes indicate respectively the first and third quartiles; the whiskers define the minima and maxima; data were analyzed using the Mann-Whitney. Survival curves were analyzed using the LogRank test.

### The Toll pathway is required in the host defense against some *A. fumigatus* tremorgenic mycotoxins

The *ftm* gene cluster of *A. fumigatus* has been shown to be involved in the biosynthesis of secondary metabolites belonging to the tremorgenic toxins such as the fumitremorgins and verruculogen. The *ftmA* gene encodes the first enzyme of this biosynthetic pathway (Kato et al., 2013). As shown in Fig. 3*A*, a *ΔftmA* mutant was slightly but reproducibly less virulent than its wild-type *A. fumigatus* control strain. Whereas *MyD88* and wild-type flies behaved similarly after the injection of either low or high doses of verruculogen, *MyD88* flies were more sensitive than wild-type to this toxin injected at a 1 or 5 mg/ml concentration (or introduced as a powder thereby bypassing the need for dissolution in a DMSO-containing solvent), in conventional or microbe-free conditions (Fig. 3*B-D*, Fig. EV5*A*). *Toll* and *spz* mutant flies also succumbed to injected verruculogen (Fig. EV5*B-C*). *MyD88* flies were also sensitive to fumitremorgin C injected at concentrations greater than or equal to 1 mg/ml (Fig. 3*E*).

**Figure 3.**
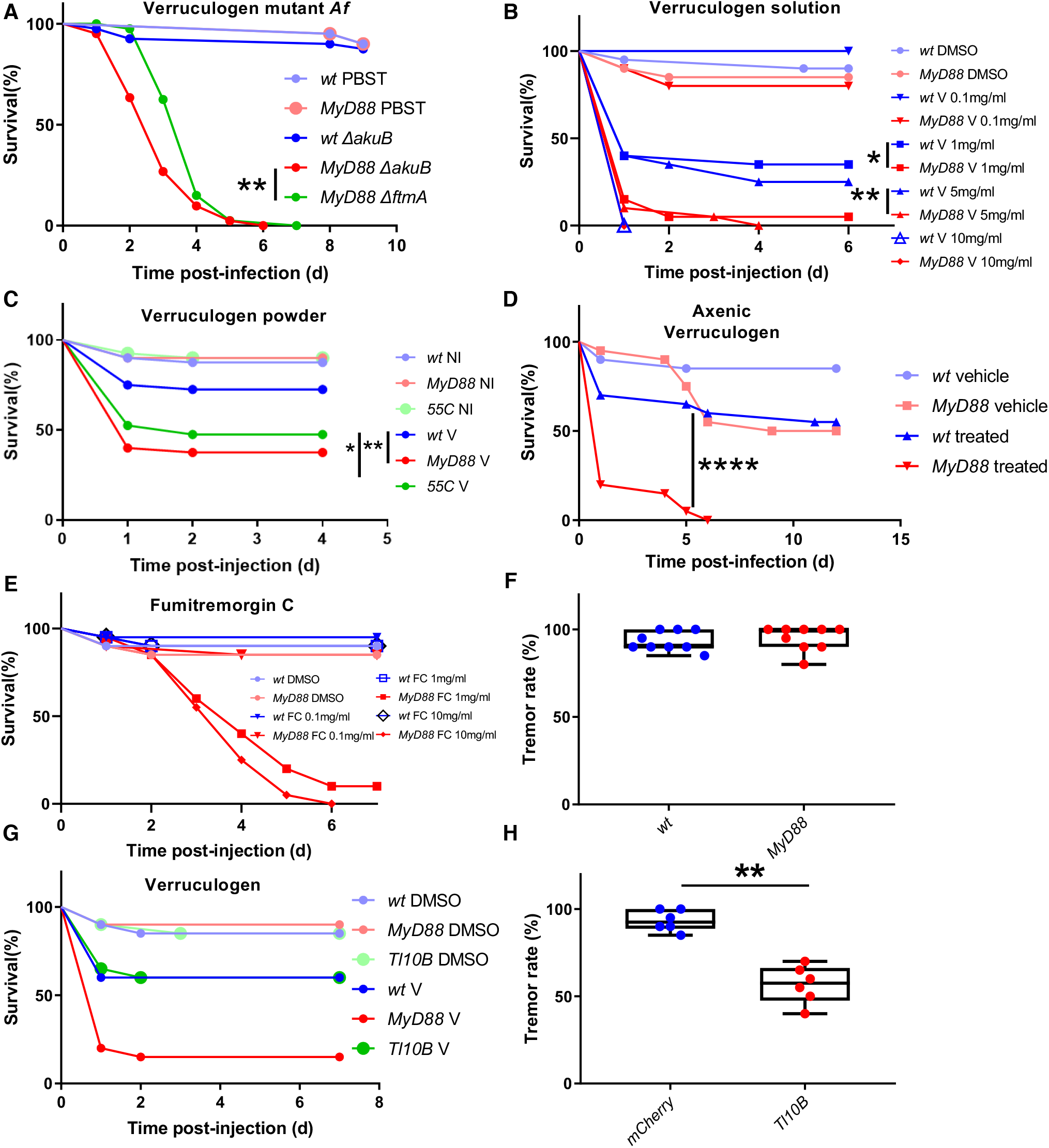
The Toll pathway mediates *Drosophila* resilience to *A. fumigatus* tremorgenic secondary metabolites of the fumitremorgin/verruculogen biosynthesis pathway. (*A-B*) survival of *MyD88* or wild-type flies after injection of 250 conidia of *ΛλftmA* (verruculogen and fumitremorgins biosynthesis pathway mutant) or wild-type [*ΛλakuB*] *A. fumigatus* (*A*) or verruculogen (V) (*B*) (n = 20 per condition); *ΛλakuB vs. ΛλftmA*, p=0.002 (*A*); wt *vs. MyD88*(1 mg/mL verruculogen, p=0.015; 5 mg/mL, p=0.008 (*B*)). (*C-D*) survival of *MyD88* mutant flies after verruculogen powder challenge (*C*), and axenic *MyD88* mutant flies after verruculogen solution injection (*D*); wt V *vs. 55C* V, p=0.02, *vs. MyD88* V, p=0.002 for verruculogen powder challenge. (*E*) survival of *MyD88* mutant flies after fumitremorgin C injection at different concentrations. (*F*) tremor rates of batches of 20 wild-type or *MyD88* flies 3h after verruculogen injection. (*G*) *Ubi-Gal4*>*UAS*-*Toll*^*10B*^ flies survive as well as wild-type flies to verruculogen injection (n=20). (*H*) Rate of *Ubi-Gal4*>*UAS*-*Toll*^*10B*^ flies exhibiting tremors 3h after injection of verruculogen in batches of 20 flies, each dot representing one batch; p=0.002. *G-H*: the middle bar of box plots represents the median and the upper and lower limits of boxes indicate respectively the first and third quartiles; the whiskers define the minima and maxima, data were analyzed using the Mann-Whitney. Survival curves were analyzed using the LogRank test. Except indicated otherwise (*B*), the concentration of injected verruculogen was 1 mg/ml.

Most wild-type and *MyD88* flies injected with verruculogen exhibited seizures as early as half an hour after injection and by three hours all flies suffered from tremors (Fig. 3*F* and Movies EV1-4). Interestingly, wild-type flies started recovering from seizures after verruculogen injection from 15 hours onward; all surviving flies had recovered after about a day whereas *MyD88* flies never recovered (Fig. EV5*D*). Of note, when challenging directly with verruculogen powder, *MyD88* flies did recover, but slower, likely because in this mode a lower effective dose of the mycotoxin is delivered (Fig. EV5E). Upon closer inspection, we found that *MyD88*, but not wild-type flies, exhibited tremors after two days of *A. fumigatus* infection (Movie EV5). The Toll pathway is constitutively activated in *Toll*^*10B*^ flies. As expected, *Toll*^*10B*^ flies survived verruculogen injection like wild-type flies (Fig. 3*G*). Remarkably, about 50% of these flies did not exhibit tremors at three hours post injection of verruculogen (Fig. 3*H*). These data indicate that wild-type flies undergo the tremorgenic action of verruculogen and, in contrast to *MyD88* flies, are able to overcome the effects of the toxin in a resilience process that involves *spz, Toll* and *MyD88*. Melanization and hemocytes did not appear to be involved in resilience to verruculogen action (Fig. EV5*F-H*).

### The Toll pathway is required in the host defense against a ribotoxin

We next tested the contribution of another mycotoxin, restrictocin, a ribotoxin protein secreted by *A. fumigatus* and other pathogenic fungi. Restrictocin cleaves 28S ribosomal RNA and thereby inhibits host cell translation (Fando et al., 1985; Lamy et al., 1991; Nayak et al., 2001). Injection of *A. fumigatus* Δ*aspf1* conidia, which lack the restrictocin biosynthesis locus, resulted in a modest but reproducible reduction in virulence as compared to a wild-type *A. fumigatus* control strain when injected into *MyD88* mutants (Fig. 4*A*). Strikingly, the injection of restrictocin killed *MyD88* but not wild-type flies in untreated, antibiotics-treated or axenic flies (Fig. 4*B-D*). Whereas the injection of restrictocin led to the demise of *spz* and *Toll* mutant flies, it did not impact flies deficient for either melanization or the cellular immune response (Fig. EV6).

**Figure 4.**
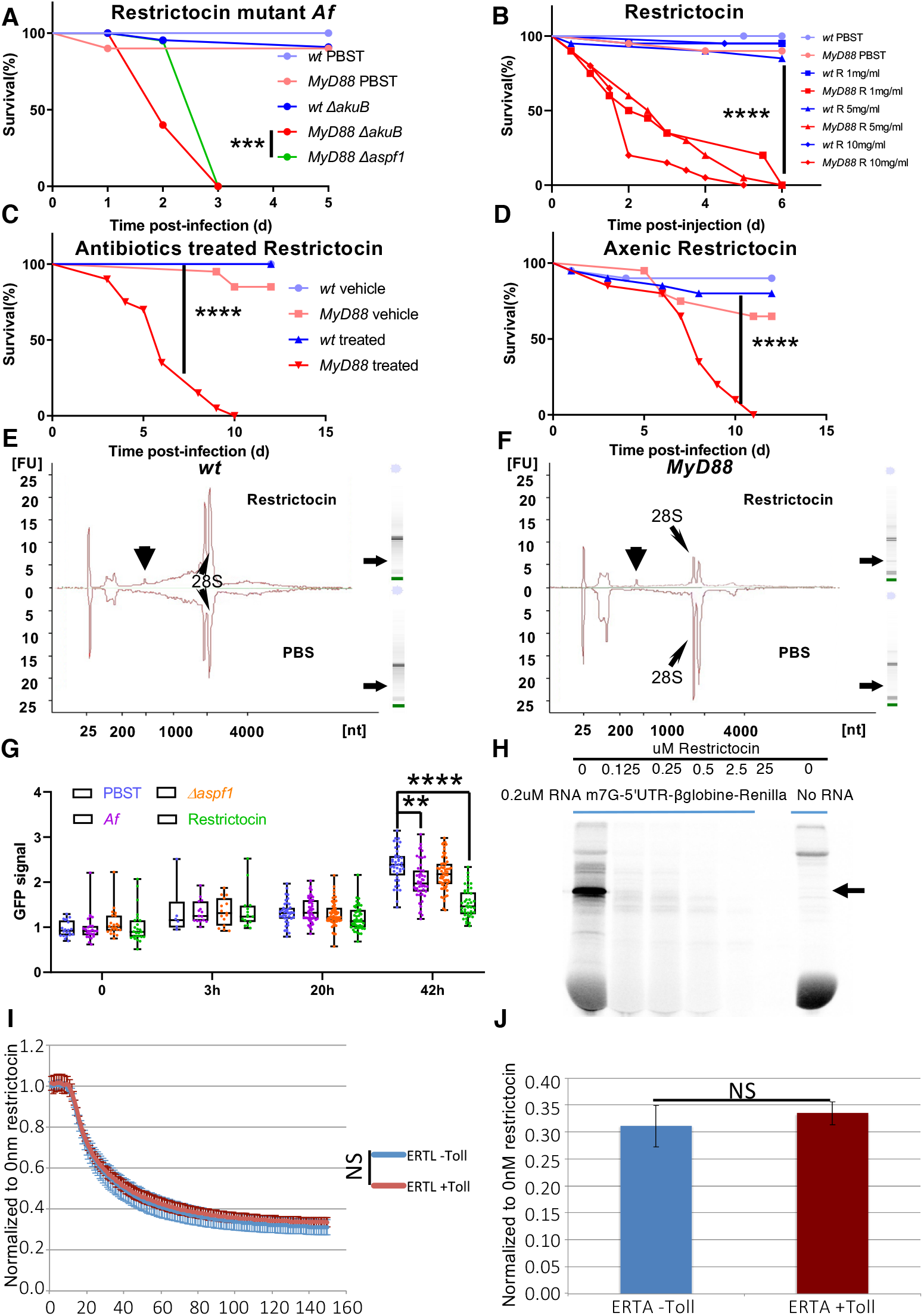
Restrictocin functions as a ribotoxin *in vitro* and *in vivo* and affects *MyD88* mutant and not wild-type flies. (*A*) survival of *MyD88* or wild-type flies to 250 injected *Δaspf1* (restrictocin mutant) or wild-type [*ΔakuB*] *A. fumigatus* conidia (n = 20); *MyD88: Δaspf1 vs. ΔakuB* p=0.0007. (*B*) survival of *MyD88* flies after the injection of different concentrations of restrictocin (R) (n = 20). (*C-D*) survival of antibiotics-treated (*C*) and axenic (*D*) *MyD88* mutant flies after restrictocin injection. (*E-F*) ribosomal RNA cleavage measurement after restrictocin or PBS injection in wild-type (*E*) and *MyD88* (*F*) flies, the arrowheads show the position of the α-sarcin fragments. (*G*) fluorescence (arbitrary units) emitted by transgenic p*Ubi-Gal4-Gal80ts*>*UAS-GFP* whole flies induced at the same time as the challenge and measured at the indicated time points; PBST *vs. Af*: p=0.002. (*H*) SDS-PAGE analysis of ^35^S*-*labelled translated proteins produced in a rabbit reticulocytes lysate from a m^7^G-capped reporter RNA containing the 5’UTR of β-globin followed by the Renilla luciferase coding sequence, in the presence of increasing concentrations (0.125 to 25 nM) of restrictocin. (*I-J*) fluorescence analysis from *in vitro* translated eGFP from an IGR (CrPV)-driven reporter in noninduced (blue) and Toll-induced (red) ERTL lysates: translation kinetics of *in vitro* synthesized eGFP in the presence of 1 nM restrictocin showing fluorescence values normalized to untreated translation reactions (*I*) and a histogram representing the end-point fluorescence quantification of I (*J*). Error bars represent the standard error of the mean from three independent experiments. *G*: the middle bar of box plots represents the median and the upper and lower limits of boxes indicate respectively the first and third quartiles; the whiskers define the minima and maxima; data were analyzed using the Kruskall-Wallis and Dunn’s post-hoc test. *J*: error bars represent the standard error of the mean from three <u>technical</u> replicates. Survival curves were analyzed using the LogRank test. NS: not significant.

The cleavage by restrictocin of the 28S RNA between G_4325_ and A_4326_ yields a fragment of about 500 nucleotides known as the α-sarcin fragment (Gluck et al., 1994). When we analyzed total RNA extracted from *MyD88* restrictocin-injected flies, we observed a fragment of the expected size, which was not detected in PBS-injected flies. The α-sarcin peak was also detected upon the injection of restrictocin in wild-type flies. The cleavage of the 28S RNA was however much less pronounced in wild-type flies as compared to *MyD88* (Fig. 4*E-F*). These observations suggest that the MyD88-mediated response is able to counteract restrictocin *in vivo* prior to its action on rRNA. In agreement with these results, the GFP fluorescence emitted from a transgene induced ubiquitously at the time of the challenge was reduced 42 hours after restrictocin injection. In addition, GFP fluorescence was lower upon *A. fumigatus* challenge than upon a mock infection (Fig. 4*G*). Taken together, these data suggest that restrictocin is able to inhibit translation to a detectable degree *in vivo*, likely through the cleavage of the ribosomal 28S α-sarcin/ricin loop as described *in vitro*.

We therefore checked in a rabbit reticulocyte translation assay that restrictocin is blocking translation *in vitro* (Fig. 4*H*), as previously reported (Nayak et al., 2001). This observation was extended to *Drosophila* S2 cell extracts. Since the Toll pathway cannot be induced in regular S2 cells, we used a stable line that expresses a chimeric Toll receptor (ERTL) that can be activated by adding Epidermal Growth Factor (EGF) to the growth medium (Sun et al., 2004). In extracts from noninduced cells, eGFP *in vitro* translation was inhibited by the addition of restrictocin in a dose-dependent manner (Fig. EV7*A-B*). Even though the Toll pathway was indeed activated by the addition of EGF, translation with an extract made from induced ERTL-S2 cells was nevertheless inhibited by the addition of restrictocin almost as efficiently as with an extract made from noninduced ERTL-S2 cells (Fig. 4*I-J*). Thus, the Toll pathway may not act at the intracellular level but possibly through secreted effectors as detailed further below.

The Spätzle/Toll/MyD88 cassette is thus required for host defense against both verruculogen, a secondary metabolite, and restrictocin, a protein ribotoxin.

### Bomanins mediate resilience to mycotoxins

The Toll pathway regulates the expression of tens of genes, including some Bomanins initially identified as *Drosophila*-immune induced molecules (Uttenweiler-Joseph et al., 1998). Strikingly, the deletion of the 55C locus (Fig. EV2A) that spans ten *Bomanin* genes yields a phenotype as strong as Toll pathway mutants in several infection models (Clemmons et al., 2015). In the case of *A. fumigatus*, we found the *Bom*^*Δ55C*^ deletion mutant to be only somewhat less susceptible to this infection than *MyD88* flies (Fig. 5*A*). The fungal burden remained low during and after the infection (Fig. 5*B*, Fig. EV8*A*). Interestingly, the *Bom*^*Δ55C*^ mutant was also sensitive to the injection of verruculogen and restrictocin (Fig. 5*C-D*; green curve). Only some 25% of *Bom*^*Δ55C*^ flies *vs*. more than 50% for isogenized wild-type survived verruculogen injection after day one. In the case of restrictocin, *Bom*^*Δ55C*^ flies succumbed to this challenge, which was not the case for control flies. To exclude the possibility of a nonspecific sensitivity of *MyD88* or of *Bom*^*Δ55C*^ flies to stress, we submitted these mutant flies as well as their isogenized controls to a variety of stresses such as heat shock at 37°C or 29°C or the injection of salt solution or H_2_O_2_ (Fig. EV8*B-E*). The injection of 4.6 nL of 8% NaCl solution or of 2% H_2_O_2_ did not reveal a differential susceptibility of the immuno-deficient flies to these challenges. In contrast, we did observe a mild susceptibility of *MyD88* but not of *Bom*^*Δ55C*^ flies to a continuous exposure to 37°C. Similar results were obtained for an exposure to 29°C with *MyD88* flies displaying an enhanced sensitivity but only after 12-15 days, that is, much later than the usual time frame of our experiments (Fig. 1*A*, Fig. 3*B*, Fig. 4*B*).

**Figure 5.**
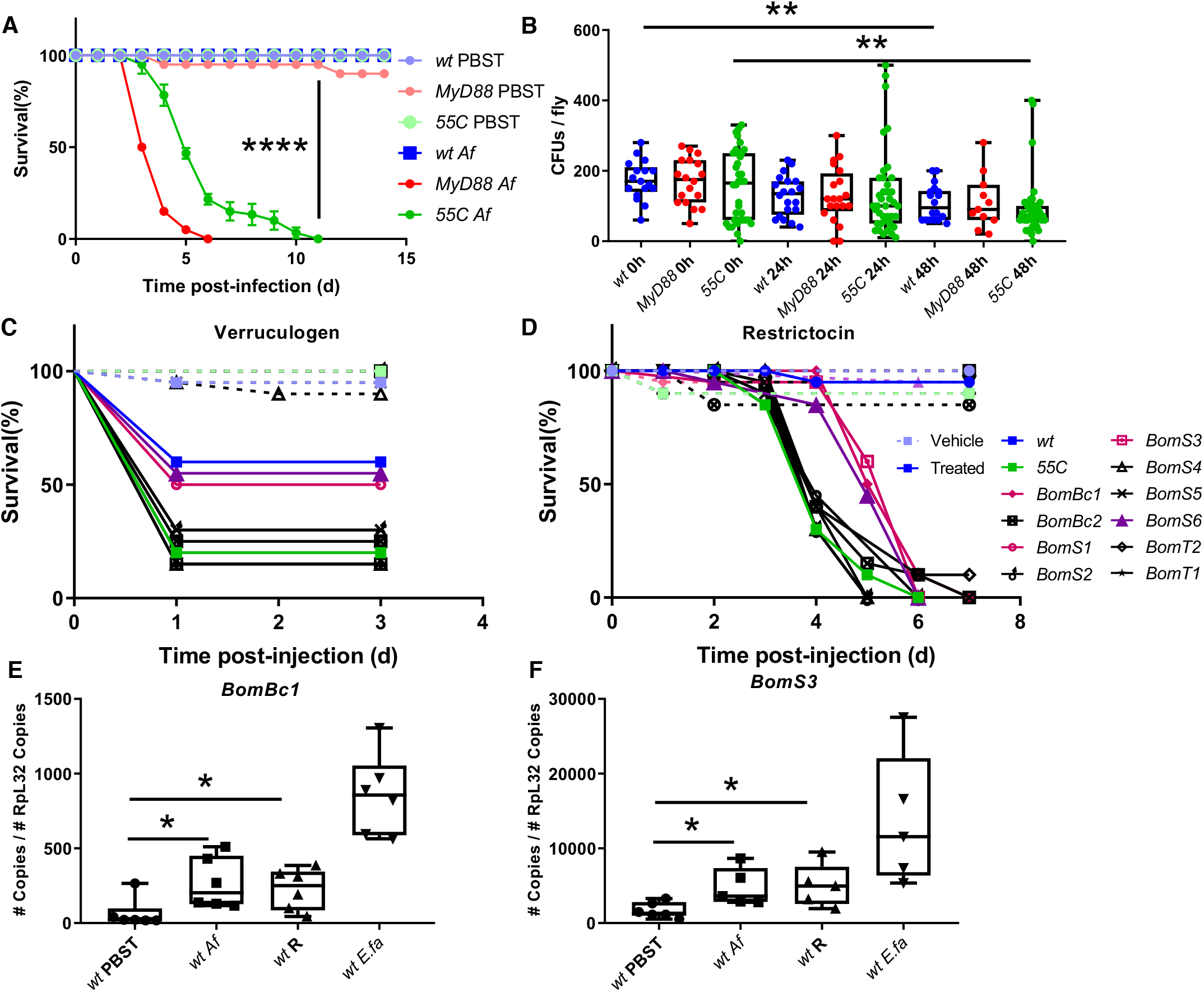
Distinct *Bomanins* mediate resilience to specific *A. fumigatus* mycotoxins. (*A-B*) survival (*A*) and fungal load (*B*) of *Bom*^*τ155C*^ (55C) deficient flies compared to wild-type and *MyD88* after injection of 250 conidia; mean and standard deviation of the survival of triplicates of 20 flies each are displayed. (*B*): the fungal burden does not increase in *Bom*^*τ155C*^-deficient flies; wt 0h *vs*. 48h, p=0.001; 55C 0h *vs*. 48h, p=0.007. (*C-D*) rescue of the sensitivity of *Bom*^*τ155C*^ flies to verruculogen (*C*) or to restrictocin (*D*) by the transgenic expression of individual 55C locus genes (indicated in the caption). 55C flies *vs. BomBc1*or *BomS3*, p<0.0001, *vs. BomS6*, p=0.0028 for restrictocin assay; and *vs. BomS1*, p=0.0495, *BomS6*, p=0.011 for verruculogen assay. (*E-F*) expression levels of *BomBc1*, and *BomS3* measured by RT-digital PCR 48 hours after challenge; *BomBc1* PBST *vs. Af*, p=0.015, PBST *vs*. restrictocin (R), p=0.015; *BomS3:* PBST *vs. Af*, p=0.02, PBST *vs*. R, p=0.03. *B,E-F*: the middle bar of box plots represents the median and the upper and lower limits of boxes indicate respectively the first and third quartiles; the whiskers define the minima and maxima; data were analyzed using the Mann-Whitney statistical test. Survival curves were analyzed using the LogRank test. NS: not significant.

We attempted to identify the relevant 55C cluster genes involved in host defense against injected mycotoxins using a genetic rescue strategy in which we overexpressed single 55C locus *Bomanin* genes in the background of the *Bom*^*Δ55C*^ deficiency. Overexpression of either *BomBc1, BomS3*, or *BomS6* provided a significant degree of protection against restrictocin whereas *BomS6* and, to a variable extent, *BomS1* protected *Bom*^*Δ55C*^ mutant flies from verruculogen and survived the challenge like wild-type flies (Fig. 5*C-D*). Of note, we still observed the induction of tremors in verruculogen-injected rescue flies. To determine whether BomS3 interacts with restrictocin *in vitro*, we tested whether the pre-incubation of restrictocin with a BomS3 synthetic peptide would decrease the inhibition of translation in ERTL-S2 cells. As shown in Fig. EV7*C-D*, it was as inefficient as control synthetic BomS1 peptide in blocking the inhibition of translation mediated by restrictocin. Thus, BomS3 is unlikely to act directly alone against restrictocin and might act extracellularly.

As monitored by RT-dPCR, the expression of *BomBc1, BomS1*, and especially *BomS3* were induced by an *A. fumigatus* challenge (Fig. 5*E-F*, Fig. EV2*D*). Some other *Bom* genes located in 55C also exhibited a weak induction, which was not significant (Fig. EV9*B-H*). As the injection of vehicle alone induced a significant response of some 55C locus genes, it was not possible to determine whether verruculogen is also able to induce their expression in this experimental series. We did find that the expressions of *BomBc1, BomS3*, and *BomS4* were induced to a low level by the injection of restrictocin (Fig. 5*E-F*, Fig. EV9*C*), which was not the case for other *Bomanin* genes (Fig. EV9*A-B,D-H*).

### BomS6 can protect flies from the toxic effects of verruculogen when expressed in the brain

That the ubiquitous overexpression of *Tl*^*10B*^ protects 50% of wild-type flies from the tremorgenic effects of verruculogen provided a convenient method to investigate which tissues mediate this effect. When we expressed the *Tl*^*10B*^ transgene in neurons, there was also a dominant protection of 50% of the flies from the tremors measured three hours after the injection of verruculogen, an effect similar to its ubiquitous expression (Fig. 3*H*, Fig. 6*A*). When we next monitored the survival of these flies to injected verruculogen, in contrast to the ubiquitous expression of *Tl*^*10B*^ in which 40% of flies succumb to this challenge, like wild-type flies (Fig. 3*G*), there was a full protection conferred to flies in which *Tl*^*10B*^ is expressed only in neurons (Fig. 6*B*). Similar observations were made when *Tl*^*10B*^ was expressed in glial cells, except that the degree of protection was weaker, possibly reflecting the possibility that Toll acts through secreted effectors (Fig. 6*C-D*). We therefore tested the ectopic expression in a wild-type background of BomS6, which is the only peptide we found to reliably rescue the sensitivity phenotype of *Bom*^*Δ55C*^ flies in survival experiments. When *BomS6* was ectopically expressed in the nervous system, all of the flies displayed tremors three hours after injection and there was no enhanced protection of this phenotype (Fig. EV10*A-B*). However, when we measured the time it took for those flies to recover from tremors, we did find that they recovered faster, at a pace similar to that obtained by its ubiquitous ectopic expression (Fig. 6*E-F*). Interestingly, flies in which *BomS6* was expressed in neurons or ubiquitously were fully protected against the noxious effects of verruculogen in survival experiments (Fig. 6*G*). When *BomS6* was expressed in glial cells, the improved recovery from verruculogen challenge was nearly significant (Fig. 6*H*, p=0.058) and flies did survive significantly better than wild-type flies, but nevertheless were less protected than when *BomS6* was expressed in neurons (Fig. 6*I*). When we tried to repeat the experiment by ectopically expressing *BomS4*, no protection was conferred to those flies suggesting a degree of specificity of the *BomS* genes (Fig. EV10*C-D*).

**Figure 6.**
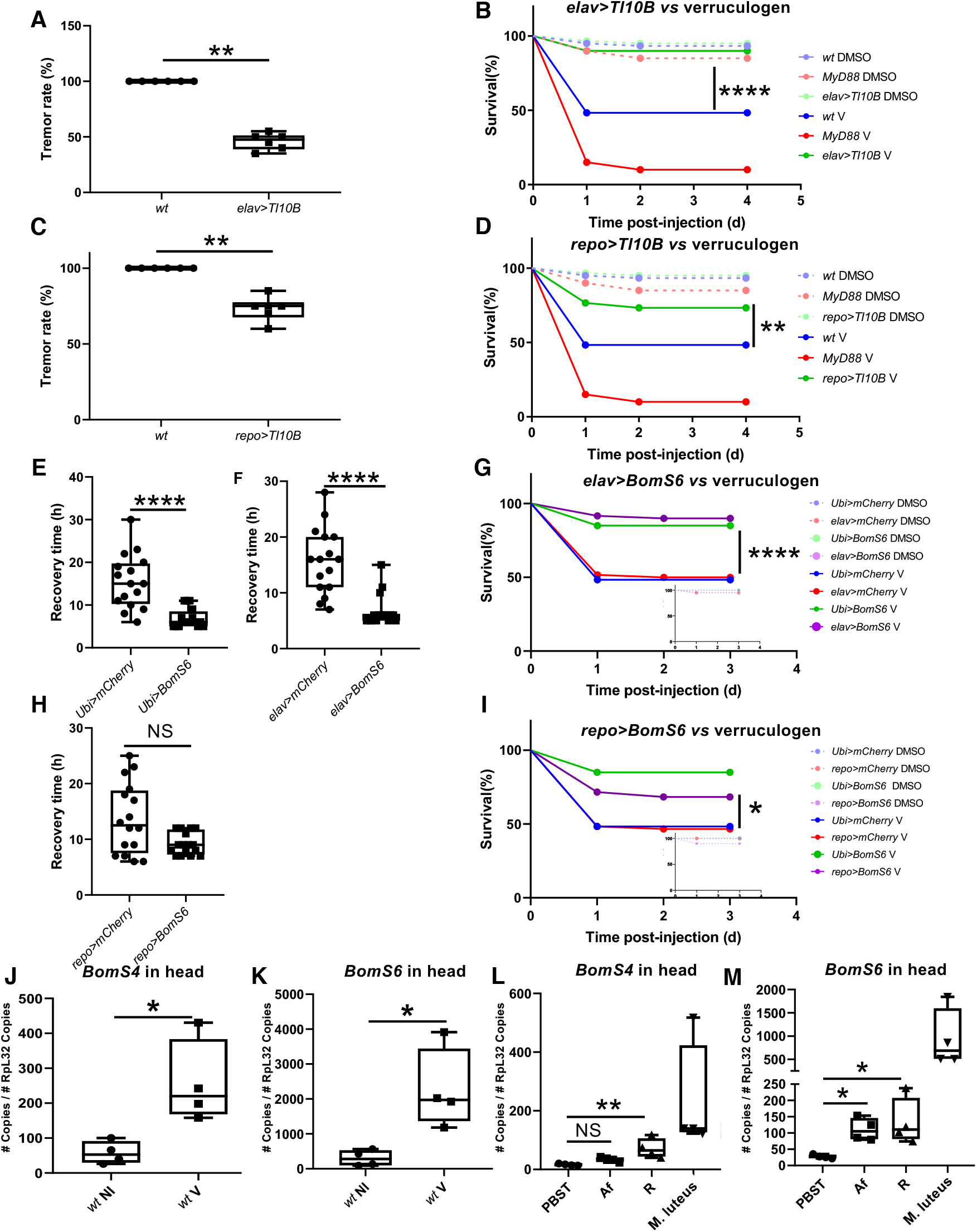
*Bomanin S6* mediates resilience to verruculogen in the nervous system of *Drosophila*. (*A-B*) tremor rate (*A*) and survival (*B*) of flies overexpressing *Tl[10B]* in neurons (n = 60) compared to wild-type after injection of verruculogen, tremor rate (*A*) *wt vs. elav> UAS*-*Toll*^*10B*^, p=0.002. (*C-D*) tremor rate (*C*) and survival (*D*) of flies overexpressing *Tl[10B]* in glia (n = 60) compared to wild-type after injection of verruculogen, tremor rate (*C*) *wt vs. repo> UAS*-*Toll*^*10B*^, p=0.002; survival (*D*) *wt* V *vs. repo> UAS*-*Toll*^*10B*^ V, p=0.005. (*E-G*) recovery time from tremor (*E,F*) and survival (*G*) of flies overexpressing *BomS6* in glia (n = 60) compared to wild-type after injection of verruculogen. (*H-I*) recovery time from tremor (*H*) and survival (*I*) of flies overexpressing *BomS6* in glia (n = 60) compared to wild-type after injection of verruculogen, recovery time (*H*) *repo>mCherry vs. repo>BomS6*, p=0.058, survival (*I*) *repo>mCherry* V *vs. repo>BomS6* V, p=0.016. (*J-M*) expression of *BomS4* (*J*) and *BomS6* (*K*) in head after verruculogen powder challenge, and *BomS4* (*L*) and *BomS6* (*M*) in head after restrictocin injection. *A,C,E,F,H,J-M*: the middle bar of box plots represents the median and the upper and lower limits of boxes indicate respectively the first and third quartiles; the whiskers define the minima and maxima; data were analyzed using the Mann-Whitney statistical test. Survival curves were analyzed using the LogRank test. NS: not significant. In this figure, concentration of injected verruculogen or restrictocin was 1 mg/ml.

To determine whether *Bomanins* can be induced by mycotoxins, we monitored their expression by dRT-PCR on head samples after verruculogen powder challenge or restrictocin injection. Strikingly, we found that only the expression of *BomS6* and *BomS4* were increased by the verruculogen powder challenge (Fig. 6*J-K*, Fig. EV11). Unexpectedly, these two genes were also the only ones to be induced in the head by the injection of restrictocin (Fig. 6*L-M*). In contrast, all 55C *Bomanin* genes were induced in the head after the injection of 500 *A. fumigatus* conidia, except for *BomS3* and *BomS4* (Fig. 6*L-M*, Fig. EV12). Of note, *Drosomycin* expression was induced in the head after an *A. fumigatus* challenge but not by restrictocin or verruculogen (Fig. EV11*I*, EV12*I*). Thus, only two *Bomanin* genes are induced in the head in response to restrictocin or verruculogen injection.

## Discussion

Here, we observed that *A. fumigatus* remains confined to its injection site in both wild-type and Toll pathway mutant flies due to the restriction of fungal dissemination, but not its annihilation, by melanization. Thus, this rare occurrence of a localized infection together with the analysis of mycotoxin mutants of *A. fumigatus* confirms the fundamental role of mycotoxins in the virulence of *A. fumigatus* and reveals an unanticipated role for the Toll pathway in the protection against various secreted poisonous molecules. In the course of evolution, host defense effectors able to effectively neutralize the action of mycotoxins have been selected independently of classical xenobiotics detoxification pathways.

### Induction of the expression of specific *Bomanin* genes upon mycotoxin challenge

The induction of *Drosomycin* transcripts by injected *A. fumigatus* conidia appears at best to be very mild, a trend already observed with the injection of the monomorphic yeast *C. glabrata*. It may be linked to the masking of ß-(1-3) glucans by hydrophobin proteins or melanin on the conidial cell wall (Blango et al., 2019; van de Veerdonk et al., 2017). The secretion of inhibitors of NF-κB signaling such as gliotoxin may also be at work (Pahl et al., 1996). It is however perplexing that secreted short Bomanins but not Drosomycin were detected by mass spectrometry in the hemolymph even though this peptide is massively produced during the systemic immune response to injected bacteria, at a concentration of 0.3 µM.

We find that the injection of restrictocin in flies leads to a modest yet significant induction of only a subset of *Bomanins, BomBc1, BomS3*, and *BomS4* (Fig. 5E-F, Fig. EV9*C*), even though all of them but *BomS4* -which has the lowest basal expression- are induced by a systemic immune challenge by *M. luteus* (Fig. EV2*B*). As regards the expression of *Bomanins* in the head by verruculogen, only *BomS4* and *BomS6* were induced (Fig. 6*J-K*, Fig. EV11). This differential expression of *Bomanin* genes upon mycotoxin injection, especially that of *BomS4* that is not induced in the systemic immune response, suggests that the observed induction is not due to peptidoglycan contaminating the mycotoxin preparations as it would have induced almost all *Bomanins*. Of note, we have likely employed higher concentrations of mycotoxins than actually released during infection. Indeed, whereas *BomS4* is induced in heads by verruculogen powder challenge, it is not induced there after *A. fumigatus* infection. Taken together, these data then suggest that a process akin to the immune surveillance of core cellular processes first described in *C. elegans* may also exist in *Drosophila*, especially as regards restrictocin that inhibits translation (Dunbar et al., 2012; McEwan et al., 2012; Melo and Ruvkun, 2012). We conclude that restrictocin and verruculogen induce a response limited to two *BomS* genes in the head, which is distinct from that induced in the framework of the Toll-dependent systemic immune response.

### Functions and specificity of Bomanins encoded at the 55C locus

The current paradigm for insect immune-induced secreted peptides is that they primarily represent AMPs (Hanson and Lemaitre, 2020; Lazzaro et al., 2020; Lin et al., 2020). This has been checked experimentally by the deletion of multiple AMP genes loci; the deletion of AMP genes regulated mostly by the IMD pathway phenocopied the susceptibility to Gram-negative bacteria of IMD pathway mutants (Hanson et al., 2019). This was however less clear as regards the deletion of AMP genes regulated by the Toll pathway. It appears that 55C locus *Bomanin* genes play a predominant role in the host defense against Gram-positive bacteria, yeasts, and fungi (Hanson et al., 2019). It is clear that some of these genes are required in the resistance against *E. faecalis*, suggesting that some Bomanins may function as AMPs, a hypothesis we have independently confirmed at the genetic level (Clemmons et al., 2015; Lou, 2021). *Bom*^*Δ55C*^ deficiency flies are susceptible to *C. glabrata* infection and this susceptibility was rescued by overexpressing *BomS* genes such as *BomS3* (Lindsay et al., 2018). However, no *C. glabrata* killing activity of synthetic Bom peptides could be found in *in vitro* assays (Lindsay et al., 2018; David Rabel, PB, unpublished data), even though hemolymph collected from wild-type but not mutant flies was fungicidal. The lack of fungicidal activity in the hemolymph of *Bom*^*Δ55C*^ flies was partially restored in the hemolymph of *Bom*^*Δ55C*^ mutant flies overexpressing *BomS5*, suggesting that at least this peptide may have some candicidal activity when combined with other Toll-dependent gene product(s) (Lindsay et al., 2018).

While that study suggested that BomS peptides are interchangeable against *C. glabrata* provided they are expressed at sufficiently high levels, we report here that the BomS peptides appear to be much more specific as regards the activity against mycotoxin action. Indeed, in the setting of the *Bom*^*Δ55C*^ deficiency, only overexpressed BomS6 or BomS1 show some activity against verruculogen, whereas the forced expression of BomS6, BomS3, or BomBc1 appears to be able to counteract restrictocin. It is worth noting that the AMP role proposed for BomS3 against *C. glabrata* by Lindsay *et al*. (2018) was based mostly on survival experiments. It thus cannot be formally excluded that BomS3 might also act against an unidentified *C. glabrata* secreted toxin. In this respect, it has recently been reported that *C. glabrata* is able to invade the brain (Benmimoun et al., 2020) where we suspect that the activation of the Toll pathway signaling takes place.

With respect to host defense against restrictocin, our partial rescue data of the *Bom*^*Δ55C*^ susceptibility phenotype by three 55C *Bomanins*, including one encoding a bicipital Bomanin might be accounted for by some form of redundancy. We cannot however exclude that these Bomanins provide a degree of protection through separate mechanisms, especially since the two Bomanin domains of Bc1 are rather divergent when compared to the high degree of conservation exhibited by BomS domains (Clemmons et al., 2015). Yet, *BomS6* is the sole 55C *Bomanin* providing protection against both restrictocin and verruculogen. BomS6 contains a lysine residue at position 10 of its Bomanin domain, like BomS2 but unlike BomS4 that contains a valine whereas other BomS peptides have an arginine at this position. The other difference is an isoleucine instead of valine at position 14 of the Bomanin domain. Thus, these biochemical differences along the capacity to be induced in the heads account for the unique function of BomS6 among Bomanins.

Whereas we propose here a specific function for some Bomanins in counteracting the effects of restrictocin or verruculogen, we cannot formally exclude an AMP function in other contexts. Indeed, it has previously been reported that mammalian alpha-defensin AMPs are also able to directly neutralize secreted bacterial virulence factors such as pore-forming toxins or enzymes that need to penetrate inside eukaryotic cells to act on their intracellular target (Kudryashova et al., 2014 and references therein). These proteins are inherently thermodynamically unstable as they need to change their conformations to insert or go through the mammalian cell plasma membrane. It has been shown that the amphipathic properties of alpha-defensins allow them to destructure these secreted virulence factors through hydrophobic interactions and thereby inactivate them (Kudryashova et al., 2014). Such a mechanism might be at play as regards a potential interaction of Bomanins with restrictocin, which does cross the plasma membrane. Indeed, the N-terminal part of mature BomS6 appears to be rather hydrophobic (38% of residues are hydrophobic) and uncharged. As regards verruculogen, it acts through hydrophobic interactions with one of its molecular targets (see further below) and possibly BomS6 might also directly interact with verruculogen through hydrophobic interactions, although other BomS peptides exhibit similar or higher hydrophobicity, for instance BomS3.

The mechanism of action of restrictocin in inhibiting translation is well established and it crosses the plasma membrane of insects easily. Thus, it may act ubiquitously on all cell types and organs of the host. Wild-type flies tolerate an exposure to a relatively large range of restrictocin concentrations whereas *MyD88* flies succumb faster to a high dose of 10mg/mL (Fig. 4*B*); the Toll-dependent response to *A. fumigatus* is not dose-dependent (Fig. EV2*G*). Toll in wild-type flies blocks to a large extent the action of restrictocin since it prevents the cleavage of 28S rRNA (Fig. 4*E-F*). BomS3 however does not appear to directly bind to restrictocin (Fig. EV7*C-D*). Taken together, these observations suggest an indirect mode of action of Bomanins, at least for BomS3 and BomS6.

Our understanding of the action(s) of verruculogen on the nervous system is less clear as multiple effects are reported in the literature. These include increased spontaneous release of glutamate and aspartate from cerebrocortical synaptosomes (Hotujac et al., 1976; Norris et al., 1980), inhibition of the GABA_A_ receptor (Gant et al., 1987a) or inhibition of calcium-activated K^+^ channels (Knaus et al., 1994) such as *Drosophila* Slowpoke, for which a detailed structural understanding of the interaction is available (Raisch et al., 2021). In contrast to the effect of restrictocin, verruculogen induces tremors also in wild-type flies, but unlike *MyD88* flies, these are able to reverse this effect, a situation also observed in cattle (Gant et al., 1987b; Norris et al., 1980). The constitutive activation of the Toll pathway neutralizes to a significant extent the tremorgenic effects of injected verruculogen early on. Interestingly, BomS6 appears to function somewhat differently when overexpressed in the nervous system of wild-type flies. It does not prevent the initial tremors induced by verruculogen but allows the host to recover more rapidly, which suggests that two distinct processes are at play. Thus, Tl^10B^ may function through an effector that is distinct from BomS6 in the early protection against tremors; alternatively, this other effector may act in concert with BomS6. Future studies should determine whether the actions of Bomanins are direct or indirect, the latter being more likely given the different targets of restrictocin, verruculogen, and fumitremorgins. One may wonder whether Bomanins might alter the permeability of the plasma membrane for instance. The recent finding of a role for another Toll pathway effector, BaramicinA, in glial cells against a neurotoxin opened the possibility of an indirect role of BaraA in regulating the permeability of the Blood-Brain-Barrier (Huang *et al*., submitted).

### Perspectives

Our work presented here as well as in a concurring study (Huang *et al*., submitted) suggests that host defense has evolved to select mechanisms not only to directly fight off invading microorganisms such as AMPs but also to protect the host against the toxins they secrete. The identification of specific effectors of *Drosophila* innate immune signaling pathways evolved to counteract toxins is an important addition to our just emerging understanding of host strategies implemented to cope with such microbial weapons, *e*.*g*., pore-forming toxins, fungal toxins in the gut, alpha-defensins (Chikina et al., 2020; Greaney et al., 2015; Kudryashova et al., 2014; Lee et al., 2016). It will be interesting to determine whether innate immunity also protects at least to some extent against mycotoxins that contaminate food that present a major health threat for animals as well as humans (Brown et al., 2021).

Aspergillosis causes acute or chronic infections in an estimated 14 million patients(Gago et al., 2019; Kosmidis and Denning, 2015). Chronic infections represent major threats to the survival of patients with comorbidities. It will therefore be important to establish whether mammalian antifungal immune response pathways also contribute to resilience against mycotoxins as is the case for the Toll pathway in flies. Finally, our findings open the possibility of the existence of host defenses that protect immunocompetent animals or humans against some mycotoxins but leave individuals deficient for these defenses susceptible to disease.

## Materials and Methods

### Microbial strains

*Aspergillus fumigatus* was cultured on potato dextrose agar (PDA) medium supplemented with 0.1g/l chloramphenicol in an incubator at 29 °C. Conidia were harvested after 4-7 days of culture. The conidial suspension was purified by filtration on cheese cloth to eliminate hyphae and other impurities. The standard wild-type *A. fumigatus CEA17ΔakuB*^*Ku80*^ (CEA17) is a kind gift from Drs. Anne Beauvais and Jean-Paul Latge (Institut Pasteur, Paris) and is also the genetic background control for *ΔgliP*. Other wild-type strains include D141 (background for D141-GFP), Af293, ATCC46645, A1160 (background for *ΔpptA*), GFP labeled strain (D141-GFP). *ΔpptA* (secondary metabolites free mutant) and *ΔgliP* (gliotoxin-free mutant) have been previously described (Hillmann et al., 2015; Johns et al., 2017).

For targeted deletion of *ftmA* (AFUB_086360) and *aspf1* (AFUB_050860) gene replacement cassettes were generated by three-fragment based PCR as described previously (Szewczyk et al., 2006). In brief, deletion constructs were generated by amplifying around 1 kb up- and downstream sequences of the respective gene and insertion of the pyrithiamine resistance cassette (Kubodera et al., 2000) by fusion PCR. Protoplasts of *CEA17ΔakuB*^*Ku80*^ *A. fumigatus* strain (da Silva Ferreira et al., 2006) were transformed with purified PCR products. Transformants were selected for resistance to pyrithiamine. Homologous recombination and integration of the deletion cassette were validated by PCR. Phusion Flash High-Fidelity Master Mix (Thermo Scientific, Germany) was used for all reactions. *A. fumigatus* was cultivated in *Aspergillus* minimal medium(Jahn et al., 1997). Media were supplemented with 0.1 mg/l pyrithiamine (Merck, Germany) when required.

The sequence of primers is found in Table EV1.

*Micrococcus luteus* CGMCC#1.2299 was cultured in Tryptic soy broth (TSB), and *E. faecalis* CGMCC#1.2135 was cultured in Luria-Bertani (LB) at 37 °C for 24h. The bacteria were then washed in PBS thrice and resuspended.

### Fly strains

Fly lines were raised on food at 25 °C with 65% humidity. For 25 l of fly food medium, 1.2 kg cornmeal (Priméal), 1.2 kg glucose (Tereos Syral), 1.5 kg yeast (Bio Springer), 90 g nipagin (VWR Chemicals) were diluted into 350 ml ethanol (Sigma-Aldrich), 120 g agar-agar (Sobigel) and water qsp were used.

*w*^*A5001*^ flies were used as wild-type control unless otherwise indicated. Canton-S (BDSC64349), *w*^*1118*^ (VDRC60000), and *y*^*1*^*w*^*1*^ were used as further wild-type controls as needed. The following mutant lines were used: *MyD88*^*c03881*^, *Df (2R)3591, Hayan*^*34*^, *w, P{ry, Dipt-LacZ}, P{w*^*+*^, *drs-GFP }; spz*^*rm7*^, *spz*^*u5*^, *Toll*^*632*^. The following strains *Df(3R)Tl-I, e*^*1*^*/TM3, Ser*^*1*^ (BDSC1911), *eater*^*1*^, *SP7, PPO1*^*Δ*^, and *PPO2*^*Δ*^ were kind gifts from Dr. Bruno Lemaitre; *eater*^*Δ*^ mutants were obtained by crossing the two deficiency lines *Df(3R)Tl-I, e*^*1*^/*TM3, Ser*^*1*^ (BDSC1911) and *Df(3R)D605/TM3, Sb*^*1*^ *Ser*^*1*^ (BDSC823). *Spz* and *Tl* mutants used in this study were either transheterozygous or hemizygous mutants crossed at 25°C. *phmlΔ*-Gal4>UAS-eGFP is a reporter line for hemocytes, *w*, P{UAS-*rpr*.C}, P{UAS-*hid*} flies (a kind gift of Akira Goto) were crossed to the *phmlΔ*-*Gal4*>*UAS-eGFP* line at 29°C to ablate hemocytes during development. *UAS-Toll*^*10B*^ flies were crossed to a *w; pUbi-Gal4, pTub-Gal80ts* (BDSC30140) driver line at 25°C; hatched adults were placed at 29°C for five days to activate the Toll pathway.

The *Bom*^*Δ55C*^ deficiency was a kind gift of Steven Wasserman that was further isogenized in the *w*^*A5001*^ background. The transgenic lines expressing single *Bom* genes of the 55C locus under the *pUAS-hsp70* promoter control were generated as described (list of primers in Table S1) and checked by sequencing. The transgenic flies were crossed to a *w; pUbi-Gal4, pTub-Gal80*^*ts*^ driver line, in a homozygous *Bom*^*Δ55C*^ mutant or *w*^*A5001*^ background. The expression of the transgenes was checked by RTqPCR and mass-spectrometry analysis on collected hemolymph of single flies as required.

### Preparation of toxin solutions

Restrictocin (Sigma) was resuspended in phosphate buffer saline (PBS) pH = 7.2, gliotoxin (Abcam), helvolic acid (Abcam), fumagillin (Abcam), verruculogen (Abcam), fumitremorgin C (Sigma), were dissolved at 10 mg/mL in Dimethyl sulfoxide (DMSO; Molecular biology grade, Sigma) as stock solutions and stored at -20°C. A working concentration of 1 mg/ml in DMSO was used for injections of 4.6 nL of all toxin solutions unless otherwise indicated. Toxin solutions were thawed on ice for one hour prior to use. As multiple freeze/thaw cycles reduce the potency of the toxins, care was taken not to use an aliquot more than five times and aliquots were not stored for more than one month.

### Axenic flies

To obtain axenic flies, eggs were collected, washed with water and then 70% ethanol prior to dechorionation of eggs in a solution of 50% bleach until the chorion disappeared. Eggs were transferred into sterile vials containing media and a mix of antibiotics: ampicillin, chloramphenicol, erythromycin and tetracycline. Once emerged, adult flies were crushed and tested on LB-, Brain-Heart infusion Broth-, MRS and yeast peptone dextrose-agar plate to observe any contamination by bacteria or fungi. Of note, no anaerobic microorganisms have been detected in the *Drosophila* microbiota.

Flies treated with antibiotics were fed on food containing ampicillin, tetracycline, chloramphenicol, erythromycin, and kanamycin at 50 μg/ml final concentration each. Females were collected after two generations on fly food with antibiotics and checked for sterility by plating.

### *A. fumigatus* infections and injection of toxins

For *A. fumigatus* infections, spores were prepared freshly for each infection. Unless otherwise stated, spores were injected into the thorax (mesopleuron) of adult flies, usually at a concentration of 250 to 500 spores in 4.6 nl PBS containing 0.01% Tween20 (PBST) unless indicated otherwise, using a microcapillary connected to a Nanoject II Auto-Nanoliter Injector (Drummond). The same volume of PBS-0.01% Tween20 (PBST) was injected for control experiments. All experiments were performed at 29 °C unless otherwise indicated. Prior to all infection experiments, the flies were incubated in tubes containing only 100 mM sucrose solution for two days to eliminate traces of antifungal preservatives added to the regular food. Toxins were injected as for *A. fumigatus* injection, except that a toxin solution was used instead of a spore suspension. As noted, verruculogen powder was directly introduced into flies as follows: the ethanol-cleaned needles were not filled but just dipped into the powder and then used to prick the flies. The procedure was reiterated for each fly. Whereas the injected quantity is not determined with accuracy, it nevertheless yielded reproducible results, which were however weaker than when injecting verruculogen initially dissolved into DMSO. Flies were kept on regular food without preservatives after injection.

### Saturation of phagocytosis

Latex bead injection was performed as previously described (Nehme et al., 2011; Quintin et al., 2013). The injected flies were placed on 100 mM sucrose solution for 48 hours prior to injections.

### Survival tests

Survival tests were usually performed using 5–7-day old flies. 20 flies per vial in biological triplicates were maintained at 25°C. The transgenic overexpression flies were transferred from 18°C to 29°C for seven days before challenge to allow the expression of Gal4, which is repressed by the Gal80^ts^ repressor at 18°C. Surviving flies were counted every day. Each experiment shown is representative of at least three independent experiments unless indicated otherwise.

### Quantification of the fungal load

The fungal burden was determined using single adult flies per condition. Single flies were transferred into arrays of 8 tubes (Starstedt) containing two 1.4-mm ceramic beads (Dominique Dutcher) in 100 μl PBS-0.01% Tween20. Single flies were smashed by shaking using a mixer mill 300 or 400 (F. Kurt Retsch GmbH & Co. KG) at a frequency of 30/min twice for 30 seconds and plated on potato dextrose agar (PDA) plates supplemented with antibiotics. After that, the plates were enclosed with Parafilm™ and cultured at 29 °C with 65% humidity. Colony-forming units were counted after 48 h. FLUD was performed as described(Duneau et al., 2017).

### Monitoring of *A. fumigatus* infection *in vivo*

Flies were sacrificed and dissected in 8-well diagnostic microscope slides (Thermo Scientific) (Carl Zeiss). was used for negative staining of *ΔpptA’s* hyphae, by adding to each well 5 μL Uvitex-2B for 30 seconds at room temperature. Flies injected by D141-GFP or *ΔpptA* were dissected and observed under an epifluorescent Zeiss axioscope microscope (Carl Zeiss) each hour after the injection.

### Preparation UV-killed *A. fumigatus*

The conidial suspension at about 10^10^ conidia/mL was plated on dried potato dextrose agar (PDA) supplemented with 0.1g/l chloramphenicol plates and exposed twice for 3h to the UV-light of a microbiology safety hood. Plates were cultured at 29 °C with 65% humidity, after 48 h to check for the absence of colonies. The dead conidia were resuspended and counted prior to injection.

### Scanning electron microscope

Whole flies were incubated in one mL of a solution of 0.1M phosphate buffer pH 7.2, glutaraldehyde 2.5%, paraformaldehyde 2.4% final at room temperature for at least one hour. The flies were embedded in resin prior to observation with a scanning electron microscope (Hitachi S 800).

### *Drosomycin* and *Bomanins* expression measurement

Expression of *Drosomycin* and *Bom* genes were measured by RT-qPCR and RT-digital PCR as described previously (Gottar et al., 2006; Madic et al., 2016). With respect to digital PCR, the results (# copies/µL) are normalized by the counts of the *Rpl32* reference gene from the same reverse-transcribed sample, as also done for regular RTqPCR. The sequences of primers are shown in Table S1.

### Restrictocin-mediated inhibition of translation

#### 28S RNA α-sarcin fragment

Total RNA of restrictocin or PBS -injected flies was extracted using Trizol reagent on samples of 2-3 flies. Samples were loaded in the RNA 6000 Nano chip (Agilent RNA 6000 Nano, 2100 electrophoresis Bioanalyzer) to detect the peak corresponding to the α-sarcin fragment.

#### *Level of protein synthesis inhibition* in vivo

The level of inhibition of protein by injected restrictocin or the infection by *A. fumigatus in vivo* was assessed by measuring GFP fluorescence in single fly extracts. *w; pUbi-Gal4, ptub-Gal80*^*ts*^ were crossed to *w; UAS-GFP* flies at 18°C. The progeny from the cross was kept at 18°C until *A. fumigatus* or restrictocin injection; the flies were then placed at 29°C thereafter and analyzed at the indicated times. Each fly was crushed in 200ul PBS solution prior to measuring the GFP fluorescence using a Varioskan 2000 fluorometer (Thermo Fisher Scientific).

#### Preparation of in vitro translation extracts from non-induced and Toll-induced ERTL cells

ERTL cells were grown 5 days at 25°C in 25mL of culture medium. For the Toll-induced ERTL cells, the culture medium was supplemented with 2.5ug/mL recombinant mouse EGF (Sigma-Aldrich) 16 hours before harvesting.

After harvesting, cells were washed two times in cold 40 mM HEPES–KOH pH 8, 100 mM potassium acetate, 1 mM magnesium acetate, 1 mM DTT solution and resuspended at a concentration of 10^9^ cells/mL in the same buffer supplemented with 1X Halt^™^ Protease Inhibitor Cocktail EDTA-free (Thermo Scientific™). Cell lysis was performed by nitrogen cavitation with a Cell Disruption Bomb (Parr Instrument Company). The lysate was cleared by centrifugations at 4°C with 10,000g, aliquoted, frozen in liquid nitrogen, and stored at −80°C. The induction of the Toll pathway was checked by monitoring the transcript levels of *Drosomycin* by RTqPCR.

#### In vitro *translation assays in rabbit reticulocyte lysate and ERTL cell lysates*

*In vitro* translation experiments in rabbit reticulocyte lysate were performed as previously described (Martin et al., 2011). *In vitro* translation experiments in ERTL-cell lysates were performed as previously described for S2-cell lysates (Gross et al., 2017). eGFP *in vitro* translation was assessed by measuring fluorescence (λex = 485nm; λem = 520nm) every minute for 150 minutes.

### Quantification of tremors and the recovery

Tremor quantification was performed after the injection of 4.6 nL of a 1 mg/mL verruculogen solution to batches of 20 *w*^*A5001*^ flies placed afterwards in an empty vial. Biological triplicates were analyzed for each of three independent experiments. The rate of tremor cases was measured in each tube 3h after the injection. Tremor phenotype are shown in videos available in the supplementary material of this article.

The tremor recovery was measured every 30 minutes: the flies that had recovered are the ones exhibiting no tremors and able to walk upwards on the sides of an empty vial. Observations were performed every 30 minutes and flies that had recovered (exhibiting no tremors and able to walk upwards on the sides of the vial) were removed. Three independent experiments were performed.

### MALDI mass spectra analysis

MALDI spectra obtained from *Drosophila* hemolymph were acquired and analyzed using FlexControl and Flex Analysis (Bruker Daltonics) software or converted for analysis with the open source mass spectrometry tool mMass (http://www.mmass.org). A sandwich sample preparation was used, which consists in a deposition of (1) 0.5 µL of a saturated solution of 4HCCA in acetone, (2) 0.6 µL of acidified hemolymph in 0.1 µL of trifuloroaceotic acid (TFA), and (3) 0.4 µL of a saturated solution of 4HCC in a solution of acetonitrile/0.1TFA (2:1). After a soft drying, spots were acquired in a linear positive mode at an attenuation maintained adjusted between 50 and 60 using a Bruker Daltonics UltraflexIII-Smartbeam instrument.

### Quantification and statistical analysis

All statistical analyses were performed using Prism 7 or Prism 8 (GraphPad Software, San Diego, CA). The Mann-Whitney and/or Kruskall-Wallis tests were used unless otherwise indicated. The Log-rank test was used to analyze survival experiments. When using parametric tests (analysis of variance (ANOVA) and t-test), a Gaussian distribution of data was checked using either D’Agostino-Pearson omnibus or Shapiro-Wilk normality tests. All experiments were performed at least three times, unless otherwise indicated. Significance values: *p < 0.05; **p < 0.01; ***p< 0.001; ****p < 0.0001.

## Acknowledgments

We thank Anne Beauvais and Jean-Paul Latge for the *A. fumigatus* strain used in this study, Won-Jae Lee, Bruno Lemaitre, Jiyong Liu, Steven Wasserman, Akira Goto, Angela Giangrande, and the Guangzhou Drosophila Resource Center for fly stocks. Stocks obtained from the Bloomington Drosophila Stock Center (NIH P40OD018537) were also used in this study. We gratefully acknowledge the contributions of Valérie Demais from Plateforme d’Imagerie *in vitro* (UPS 3156-Université de Strasbourg) for scanning electron microscopy, Sébastien Voisin from the BioPark for MALDI-TOF analysis, and Miriam Yamba for expert technical help. We thank Adrian Acker for the gift of the ERTL S2 cells, controls, as well as advice on the *in vitro* experimental conditions. Finally, we are indebted to Matthew Blango for critical reading of the manuscript. RX and YL were respectively partially funded through the Sino-Foreign cooperative graduate education project of Guangzhou Medical University and the International Training Paln for young outstanding scientific research talents of Guangdong Province. This work was supported by the Deutsche Forschungsgemeinschaft collaborative research center/transregion 124 FungiNet (project A1) and the excellence cluster Balance of the Microverse to TH and AB, the Association Platform BioPark of Archamps on its Research & Development budget (PB), by grants from ‘Agence Nationale pour la Recherche’ (ANR-17-CE12-0025) to AT and FM, from the 111 Project (#D18010; China), the Incubation Project for Innovative Teams of the Guangzhou Medical University, the Open Project from State Key Laboratory of Respiratory Diseases, China, and the China High-end Foreign Talent Program to DF.

## Author Contributions

RX, YL, ZL, SL, and DF designed, performed and analyzed experiments, AT and FM designed performed and analyzed the *in vitro* translation experiments, YL and PB performed and analyzed the mass spectrometry analysis, TH and AB generated the *A. fumigatus* mutants reported in this study, RX, YL, and DF wrote the manuscript with inputs from all authors.

## References

Alarco, A.M., Marcil, A., Chen, J., Suter, B., Thomas, D., and Whiteway, M. (2004). Immune- deficient Drosophila melanogaster: a model for the innate immune response to human fungal pathogens. J Immunol 172, 5622–5628.

Apidianakis, Y., Rahme, L.G., Heitman, J., Ausubel, F.M., Calderwood, S.B., and Mylonakis, E. (2004). Challenge of Drosophila melanogaster with Cryptococcus neoformans and role of the innate immune response. Eukaryot Cell 3, 413–419.

Benmimoun, B., Papastefanaki, F., Perichon, B., Segklia, K., Roby, N., Miriagou, V., Schmitt, C., Dramsi, S., Matsas, R., and Speder, P. (2020). An original infection model identifies host lipoprotein import as a route for blood-brain barrier crossing. Nature communications 11, 6106.

Blango, M.G., Kniemeyer, O., and Brakhage, A.A. (2019). Conidial surface proteins at the interface of fungal infections. PLoS Pathog 15, e1007939.

Bongomin, F., Gago, S., Oladele, R.O., and Denning, D.W. (2017). Global and Multi-National Prevalence of Fungal Diseases-Estimate Precision. J Fungi (Basel) 3, 57.

Brown, R., Priest, E., Naglik, J.R., and Richardson, J.P. (2021). Fungal Toxins and Host Immune Responses. Frontiers in microbiology 12, 643639.

Chikina, A.S., Nadalin, F., Maurin, M., San-Roman, M., Thomas-Bonafos, T., Li, X.V., Lameiras, S., Baulande, S., Henri, S., Malissen, B., et al. (2020). Macrophages Maintain Epithelium Integrity by Limiting Fungal Product Absorption. Cell 183, 411–428 e416.

Clemmons, A.W., Lindsay, S.A., and Wasserman, S.A. (2015). An effector Peptide family required for Drosophila toll-mediated immunity. PLoS Pathog 11, e1004876.

Cohen, L.B., Lindsay, S.A., Xu, Y., Lin, S.J.H., and Wasserman, S.A. (2020). The Daisho Peptides Mediate Drosophila Defense Against a Subset of Filamentous Fungi. Front Immunol 11, 9.

Cramer, R.A., Jr., Gamcsik, M.P., Brooking, R.M., Najvar, L.K., Kirkpatrick, W.R., Patterson, T.F., Balibar, C.J., Graybill, J.R., Perfect, J.R., Abraham, S.N., et al. (2006). Disruption of a nonribosomal peptide synthetase in Aspergillus fumigatus eliminates gliotoxin production. Eukaryot Cell 5, 972–980.

da Silva Ferreira, M.E., Kress, M.R., Savoldi, M., Goldman, M.H., Hartl, A., Heinekamp, T., Brakhage, A.A., and Goldman, G.H. (2006). The akuB(KU80) mutant deficient for nonhomologous end joining is a powerful tool for analyzing pathogenicity in Aspergillus fumigatus. Eukaryot Cell 5, 207–211.

De Gregorio, E., Spellman, P.T., Tzou, P., Rubin, G.M., and Lemaitre, B. (2002). The Toll and Imd pathways are the major regulators of the immune response in Drosophila. Embo J 21, 2568–2579.

Dudzic, J.P., Hanson, M.A., Iatsenko, I., Kondo, S., and Lemaitre, B. (2019). More Than Black or White: Melanization and Toll Share Regulatory Serine Proteases in Drosophila. Cell reports 27, 1050–1061 e1053.

Dunbar, T.L., Yan, Z., Balla, K.M., Smelkinson, M.G., and Troemel, E.R. (2012). C. elegans detects pathogen-induced translational inhibition to activate immune signaling. Cell Host Microbe 11, 375–386.

Duneau, D., Ferdy, J.B., Revah, J., Kondolf, H., Ortiz, G.A., Lazzaro, B.P., and Buchon, N. (2017). Stochastic variation in the initial phase of bacterial infection predicts the probability of survival in D. melanogaster. eLife 6.

Fando, J.L., Alaba, I., Escarmis, C., Fernandez-Luna, J.L., Mendez, E., and Salinas, M. (1985). The mode of action of restrictocin and mitogillin on eukaryotic ribosomes. Inhibition of brain protein synthesis, cleavage and sequence of the ribosomal RNA fragment. Eur J Biochem 149, 29–34.

Fehlbaum, P., Bulet, P., Michaut, L., Lagueux, M., Brockaert, W.F., Hétru, C., and Hoffmann, J.A. (1995). Septic injury of Drosophila induces the synthesis of a potent antifungal peptide with sequence homology to plant antifungal peptides. J Biol Chem 269, 33159–33163.

Ferrandon, D. (2013). The complementary facets of epithelial host defenses in the genetic model organism Drosophila melanogaster: from resistance to resilience. Curr Opin Immunol 25, 59–70.

Frisvad, J.C., Rank, C., Nielsen, K.F., and Larsen, T.O. (2009). Metabolomics of Aspergillus fumigatus. Med Mycol 47 Suppl 1, S53–71.

Gago, S., Denning, D.W., and Bowyer, P. (2019). Pathophysiological aspects of Aspergillus colonization in disease. Med Mycol 57, S219–S227.

Gant, D.B., Cole, R.J., Valdes, J.J., Eldefrawi, M.E., and Eldefrawi, A.T. (1987a). Action of tremorgenic mycotoxins on GABAA receptor. Life Sci 41, 2207–2214.

Gant, D.B., Cole, R.J., Valdes, J.J., Eldefrawi, M.E., and Eldefrawi, A.T. (1987b). Action of tremorgenic mycotoxins on GABAA receptor. Life Sciences 41, 2207–2214.

Gao, Q., Jin, K., Ying, S.H., Zhang, Y., Xiao, G., Shang, Y., Duan, Z., Hu, X., Xie, X.Q., Zhou, G., et al. (2011). Genome sequencing and comparative transcriptomics of the model entomopathogenic fungi Metarhizium anisopliae and M. acridum. PLoS Genet 7, e1001264.

Gluck, A., Endo, Y., and Wool, I.G. (1994). The ribosomal RNA identity elements for ricin and for alpha-sarcin: mutations in the putative CG pair that closes a GAGA tetraloop. Nucleic Acids Res 22, 321–324.

Gottar, M., Gobert, V., Matskevich, A.A., Reichhart, J.M., Wang, C., Butt, T.M., Belvin, M., Hoffmann, J.A., and Ferrandon, D. (2006). Dual Detection of Fungal Infections in Drosophila via Recognition of Glucans and Sensing of Virulence Factors. Cell 127, 1425–1437.

Greaney, A.J., Leppla, S.H., and Moayeri, M. (2015). Bacterial Exotoxins and the Inflammasome. Front Immunol 6, 570.

Gross, L., Vicens, Q., Einhorn, E., Noireterre, A., Schaeffer, L., Kuhn, L., Imler, J.L., Eriani, G., Meignin, C., and Martin, F. (2017). The IRES 5’UTR of the dicistrovirus cricket paralysis virus is a type III IRES containing an essential pseudoknot structure. Nucleic Acids Res 45, 8993–9004.

Hanson, M.A., Dostalova, A., Ceroni, C., Poidevin, M., Kondo, S., and Lemaitre, B. (2019). Synergy and remarkable specificity of antimicrobial peptides in vivo using a systematic knockout approach. eLife 8, e44341.

Hanson, M.A., and Lemaitre, B. (2020). New insights on Drosophila antimicrobial peptide function in host defense and beyond. Curr Opin Immunol 62, 22–30.

Hillmann, F., Novohradska, S., Mattern, D.J., Forberger, T., Heinekamp, T., Westermann, M., Winckler, T., and Brakhage, A.A. (2015). Virulence determinants of the human pathogenic fungus Aspergillus fumigatus protect against soil amoeba predation. Environ Microbiol 17, 2858–2869.

Hotujac, L., Muftic, R.H., and Filipovic, N. (1976). Verruculogen: a new substance for decreasing of GABA levels in CNS. Pharmacology 14, 297–300.

Jahn, B., Koch, A., Schmidt, A., Wanner, G., Gehringer, H., Bhakdi, S., and Brakhage, A.A. (1997). Isolation and characterization of a pigmentless-conidium mutant of Aspergillus fumigatus with altered conidial surface and reduced virulence. Infect Immun 65, 5110–5117.

Johns, A., Scharf, D.H., Gsaller, F., Schmidt, H., Heinekamp, T., Strassburger, M., Oliver, J.D., Birch, M., Beckmann, N., Dobb, K.S., et al. (2017). A Nonredundant Phosphopantetheinyl Transferase, PptA, Is a Novel Antifungal Target That Directs Secondary Metabolite, Siderophore, and Lysine Biosynthesis in Aspergillus fumigatus and Is Critical for Pathogenicity. mBio 8, e01504–01516.

Kato, N., Suzuki, H., Okumura, H., Takahashi, S., and Osada, H. (2013). A point mutation in ftmD blocks the fumitremorgin biosynthetic pathway in Aspergillus fumigatus strain Af293. Bioscience, biotechnology, and biochemistry 77, 1061–1067.

Knaus, H.G., McManus, O.B., Lee, S.H., Schmalhofer, W.A., Garcia-Calvo, M., Helms, L.M., Sanchez, M., Giangiacomo, K., Reuben, J.P., Smith, A.B., 3rd, et al. (1994). Tremorgenic indole alkaloids potently inhibit smooth muscle high-conductance calcium-activated potassium channels. Biochemistry 33, 5819–5828.

Konig, S., Pace, S., Pein, H., Heinekamp, T., Kramer, J., Romp, E., Strassburger, M., Troisi, F., Proschak, A., Dworschak, J., et al. (2019). Gliotoxin from Aspergillus fumigatus Abrogates Leukotriene B4 Formation through Inhibition of Leukotriene A4 Hydrolase. Cell Chem Biol 26, 524–534 e525.

Kosmidis, C., and Denning, D.W. (2015). The clinical spectrum of pulmonary aspergillosis. Thorax 70, 270–277.

Kubodera, T., Yamashita, N., and Nishimura, A. (2000). Pyrithiamine resistance gene (ptrA) of Aspergillus oryzae: cloning, characterization and application as a dominant selectable marker for transformation. Bioscience, biotechnology, and biochemistry 64, 1416–1421.

Kudryashova, E., Quintyn, R., Seveau, S., Lu, W., Wysocki, V.H., and Kudryashov, D.S. (2014). Human defensins facilitate local unfolding of thermodynamically unstable regions of bacterial protein toxins. Immunity 41, 709–721.

Kupfahl, C., Heinekamp, T., Geginat, G., Ruppert, T., Hartl, A., Hof, H., and Brakhage, A.A. (2006). Deletion of the gliP gene of Aspergillus fumigatus results in loss of gliotoxin production but has no effect on virulence of the fungus in a low-dose mouse infection model. Mol Microbiol 62, 292–302.

Lamy, B., Moutaouakil, M., Latge, J.P., and Davies, J. (1991). Secretion of a potential virulence factor, a fungal ribonucleotoxin, during human aspergillosis infections. Mol Microbiol 5, 1811–1815.

Lazzaro, B.P., Zasloff, M., and Rolff, J. (2020). Antimicrobial peptides: Application informed by evolution. Science 368, eaau5480.

Lebrigand, K., He, L.D., Thakur, N., Arguel, M.J., Polanowska, J., Henrissat, B., Record, E., Magdelenat, G., Barbe, V., Raffaele, S., et al. (2016). Comparative Genomic Analysis of Drechmeria coniospora Reveals Core and Specific Genetic Requirements for Fungal Endoparasitism of Nematodes. PLoS Genet 12, e1006017.

Lee, K.Z., Lestradet, M., Socha, C., Schirmeier, S., Schmitz, A., Spenle, C., Lefebvre, O., Keime, C., Yamba, W.M., Bou Aoun, R., et al. (2016). Enterocyte Purge and Rapid Recovery Is a Resilience Reaction of the Gut Epithelium to Pore-Forming Toxin Attack. Cell Host Microbe 20, 716–730.

Lemaitre, B., and Hoffmann, J. (2007). The Host Defense of Drosophila melanogaster. Annu Rev Immunol 25, 697–743.

Lemaitre, B., Nicolas, E., Michaut, L., Reichhart, J.M., and Hoffmann, J.A. (1996). The dorsoventral regulatory gene cassette spätzle/Toll/cactus controls the potent antifungal response in Drosophila adults. Cell 86, 973–983.

Levashina, E.A., Ohresser, S., Bulet, P., Reichhart, J.-M., Hétru, C., and Hoffmann, J.A. (1995). Metchnikowin, a novel immune-inducible proline-rich peptide from Drosophila with antibacterial and antifungal properties. Eur J Biochem 233, 694–700.

Liegeois, S., and Ferrandon, D. (2022). Sensing microbial infections in the Drosophila melanogaster genetic model organism. Immunogenetics 74, 35–62.

Lin, S.J.H., Cohen, L.B., and Wasserman, S.A. (2020). Effector specificity and function in Drosophila innate immunity: Getting AMPed and dropping Boms. PLoS Pathog 16, e1008480.

Lindsay, S.A., Lin, S.J.H., and Wasserman, S.A. (2018). Short-Form Bomanins Mediate Humoral Immunity in Drosophila. J Innate Immun 10, 306–314.

Lou, Y. (2021). Study of the function of Bomanin genes at the 55C locus in Drosophila melanogaster host defense against microbial infections (Université de Strasbourg).

Macheleidt, J., Mattern, D.J., Fischer, J., Netzker, T., Weber, J., Schroeckh, V., Valiante, V., and Brakhage, A.A. (2016). Regulation and Role of Fungal Secondary Metabolites. Annu Rev Genet 50, 371–392.

Madic, J., Zocevic, A., Senlis, V., Fradet, E., Andre, B., Muller, S., Dangla, R., and Droniou, M.E. (2016). Three-color crystal digital PCR. Biomol Detect Quantif 10, 34–46.

Martin, F., Barends, S., Jaeger, S., Schaeffer, L., Prongidi-Fix, L., and Eriani, G. (2011). Cap-assisted internal initiation of translation of histone H4. Mol Cell 41, 197–209.

McEwan, D.L., Kirienko, N.V., and Ausubel, F.M. (2012). Host translational inhibition by Pseudomonas aeruginosa Exotoxin A Triggers an immune response in Caenorhabditis elegans. Cell Host Microbe 11, 364–374.

Medzhitov, R., Schneider, D.S., and Soares, M.P. (2012). Disease tolerance as a defense strategy. Science 335, 936–941.

Melo, J.A., and Ruvkun, G. (2012). Inactivation of conserved C. elegans genes engages pathogen- and xenobiotic-associated defenses. Cell 149, 452–466.

Nam, H.J., Jang, I.H., You, H., Lee, K.A., and Lee, W.J. (2012). Genetic evidence of a redox-dependent systemic wound response via Hayan protease-phenoloxidase system in Drosophila. Embo J 31, 1253–1265.

Nayak, S.K., Bagga, S., Gaur, D., Nair, D.T., Salunke, D.M., and Batra, J.K. (2001). Mechanism of specific target recognition and RNA hydrolysis by ribonucleolytic toxin restrictocin. Biochemistry 40, 9115–9124.

Nehme, N.T., Quintin, J., Cho, J.H., Lee, J., Lafarge, M.C., Kocks, C., and Ferrandon, D. (2011). Relative roles of the cellular and humoral responses in the Drosophila host defense against three Gram-positive bacterial infections. PLoS One 6, e14743.

Norris, P.J., Smith, C.C., De Belleroche, J., Bradford, H.F., Mantle, P.G., Thomas, A.J., and Penny, R.H. (1980). Actions of tremorgenic fungal toxins on neurotransmitter release. J Neurochem 34, 33–42.

Pahl, H.L., Krauss, B., Schulze-Osthoff, K., Decker, T., Traenckner, E.B., Vogt, M., Myers, C., Parks, T., Warring, P., Muhlbacher, A., et al. (1996). The immunosuppressive fungal metabolite gliotoxin specifically inhibits transcription factor NF-kappaB. J Exp Med 183, 1829–1840.

Quintin, J., Asmar, J., Matskevich, A.A., Lafarge, M.C., and Ferrandon, D. (2013). The Drosophila Toll Pathway Controls but Does Not Clear Candida glabrata Infections. J Immunol 190, 2818–2827.

Raffa, N., and Keller, N.P. (2019). A call to arms: Mustering secondary metabolites for success and survival of an opportunistic pathogen. PLoS Pathog 15, e1007606.

Raisch, T., Brockmann, A., Ebbinghaus-Kintscher, U., Freigang, J., Gutbrod, O., Kubicek, J., Maertens, B., Hofnagel, O., and Raunser, S. (2021). Small molecule modulation of the Drosophila Slo channel elucidated by cryo-EM. Nature communications 12, 7164.

Rodrigues, M.L., and Nosanchuk, J.D. (2020). Fungal diseases as neglected pathogens: A wake-up call to public health officials. PLoS Negl Trop Dis 14, e0007964.

Scharf, D.H., Heinekamp, T., and Brakhage, A.A. (2014). Human and plant fungal pathogens: the role of secondary metabolites. PLoS Pathog 10, e1003859.

Soares, M.P., Teixeira, L., and Moita, L.F. (2017). Disease tolerance and immunity in host protection against infection. Nat Rev Immunol 17, 83–96.

Sun, H., Towb, P., Chiem, D.N., Foster, B.A., and Wasserman, S.A. (2004). Regulated assembly of the Toll signaling complex drives Drosophila dorsoventral patterning. Embo J 23, 100–110.

Szewczyk, E., Nayak, T., Oakley, C.E., Edgerton, H., Xiong, Y., Taheri-Talesh, N., Osmani, S.A., and Oakley, B.R. (2006). Fusion PCR and gene targeting in Aspergillus nidulans. Nat Protoc 1, 3111–3120.

Uttenweiler-Joseph, S., Moniatte, M., Lagueux, M., Van Dorsselaer, A., Hoffmann, J.A., and Bulet, P. (1998). Differential display of peptides induced during the immune response of Drosophila: a matrix-assisted laser desorption ionization time-of-flight mass spectrometry study. Proc Natl Acad Sci U S A 95, 11342–11347.

van de Veerdonk, F.L., Gresnigt, M.S., Romani, L., Netea, M.G., and Latge, J.P. (2017). Aspergillus fumigatus morphology and dynamic host interactions. Nat Rev Microbiol 15, 661–674.

